# *Ets2* frame-shift mutant models express in-frame mRNA by exon skipping that complements Ets2 function in the skin

**DOI:** 10.1101/2020.09.27.316075

**Authors:** Yuki Kishimoto, Iori Nishiura, Shunsuke Yuri, Nami Yamamoto, Masahito Ikawa, Ayako Isotani

## Abstract

The Ets2 transcription factor has been implicated in various biological processes. An *Ets2* mutant model, which lacks the DNA-binding domain (ETS domain), was previously reported to exhibit embryonic lethality caused by a trophoblast abnormality. This phenotype could be rescued by tetraploid complementation, resulting in pups with wavy hair.

Here, we generated new *Ets2* mutant models with deletions in exon 8 and with frame-shift mutations using the CRISPR/Cas9 method. Homozygous mutants could not be obtained by natural mating as previously reported. After rescuing with tetraploid complementation, homozygous mutant mice were generated, but these mice did not exhibit wavy hair phenotype. Our newly generated mice exhibited exon 8 skipping, which led to in-frame mutant mRNA expression in the skin and thymus but not in E7.5 embryos. As this in-frame mutation contained the ETS domain, the exon 8-skipped *Ets2* mRNA was likely translated into protein in the skin that complemented the Ets2 function. Thus, these *Ets2* mutant models, depending on the cell types, exhibited novel phenotypes due to exon skipping and are expected to be useful in several fields of research.

**Summary statement:** New Ets2 mutant models showed embryonic lethal phenotype by a placental abnormality but did not exhibit a wavy hair phenotype as a previous model.

## Introduction

E26 avian leukemia oncogene 2, 3’ domain (Ets2), a member of the ETS family, is a transcription factor that contains an ETS winged helix-loop-helix DNA-binding domain (ETS domain) that binds to GGA(A/T) DNA sequences. It is conserved in various species, including mice and humans (Karim et al., 1990; Seidel and Graves, 2002; Sharrocks, 2001). Ets2 has been implicated in various biological contexts, including placentation, hair formation, mammary tumors, inflammatory responses, angiogenesis, and the pulmonary fibrosis (Baran et al., 2011; Man et al., 2003; Wei et al., 2004; Wei et al., 2009; Yamamoto et al., 1998).

In a previous study, Ets2-deficient mice (*Ets2^db1/db1^* mice), which lack the ETS domain through deletion of exons 9 and 10, were found to exhibit early embryonic lethality due to a trophectoderm abnormality. The tetraploid complementation technique could rescue this placental abnormality, allowing for survival of the offspring (Yamamoto et al., 1998), indicating that Ets2 is essential for placental development. *Ets2^db1/db1^* mice created using the tetraploid complementation technique exhibit a variety of phenotypes, such as wavy hair, curly whisker, and a rounded forehead, allowing them to be identified. However, their fertility is normal, and they exhibit no lethal phenotype after birth. Therefore, the *Ets2^db1/db1^* mouse is a useful model for studying treatment methods for placental abnormalities (Okada et al., 2007).

The generation of gene-deficient animal models is now commonly performed using CRISPR/Cas9-based genome engineering (Cong et al., 2013; Mali et al., 2013). Model organisms made using this technique can completely mimic the genome mutations found in human diseases, such as indel mutations and substitutions, which were previously difficult to generate using the conventional knockout method. Further, homozygous mutant mice can be obtained efficiently in the founder generation by directly delivering the crRNA/tracrRNA/Cas9 ribonucleoprotein complex into a mouse zygote via electroporation (Hashimoto et al., 2016). Unfortunately, if the homozygous mutant exhibits embryonic lethality, it cannot be obtained in this way. However, it is possible to obtain placental-deficient mutant mice, such as Ets2, in the founder generation using the tetraploid complementation method (Nagy et al., 1993) in combination with genome-edited zygotes or their embryonic stem cells.

Using the above strategy, we established new *Ets2* mutant mouse lines that contain a frame-shift deletion in exon 8, which is located before the ETS domain encoded by exons 9 and 10. These genomic mutations were predicted to produce a transcriptional product that would undergo nonsense-mediated mRNA decay (NMD) or, if translated, a protein lacking the ETS domain. We found that some of the phenotypes exhibited by these mice differed from the previous study, whose origin was investigated in this work.

## Results

### Generation of new Ets2 mutant mice

On the basis of a previous study (Yamamoto et al., 1998), we designed three gRNA targeted to sites in exon 8 that would induce a frame-shift mutation, leading to a deficiency in the ETS domain, encoded by exons 9 and 10 (Fig. 1A). The riboprotein complex, which consisted of three designed crRNAs, tracrRNAs, and Cas9 protein, was electroporated into one-cell stage zygotes, which developed until the eight-cell stage. These genome-edited embryos were used for the tetraploid complementation method in order to obtain homozygous mutant mice in the founder generation (Fig. 1B). However, no homozygous mutant mice were born from the 29 transferred embryos.

**Fig. 1.**
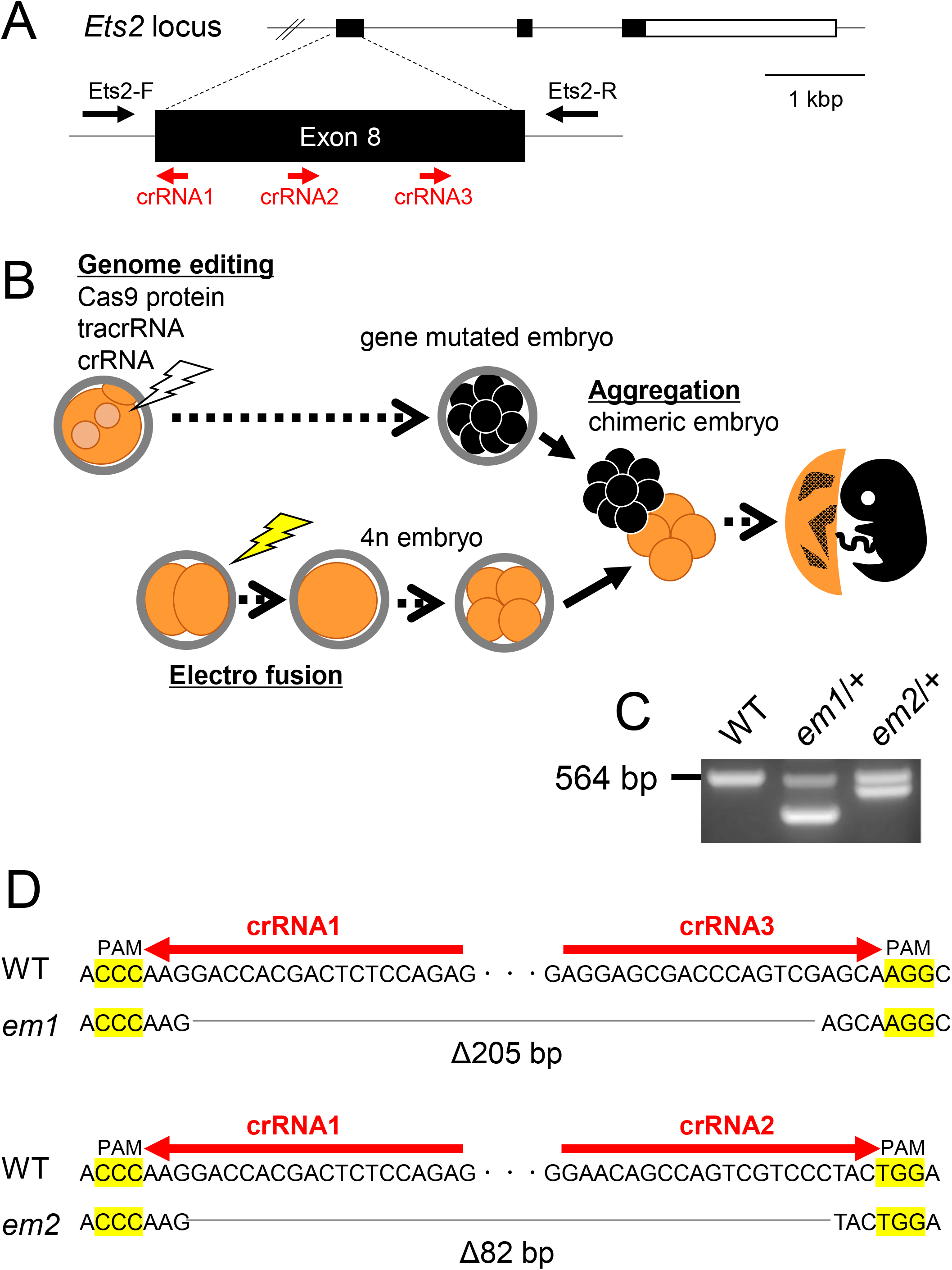
Generation of new Ets2 mutant models. (A) Design of crRNA targeting sites in exon 8 of the *Ets2* gene and checking primer positions. (B) Strategy for obtaining *Ets2* homozygous mutant 1 in F0 generation using the electroporation technique and the tetraploid method. (C) Genotyping of the two newly generated *Ets2* mutants in F0 generation. Both had a wildtype (WT: 564 bp) allele and a deletion allele (*em1* or *em2*), which were detected as shorter bands than WT. D, DNA sequence of mutant alleles and crRNA targeted sequences. The 205 bp deleted (Δ205 bp) allele was named *em1*, and the 82 bp deleted (Δ82 bp) allele was named *em2*.

Two out of three delivered pups had a heterozygous deletion mutation, which was determined by PCR analysis. One mutation was a 205-bp deletion (hereafter referred to as *em1*), and the other was an 82 bp deletion (hereafter referred to as *em2*) (Figs. 1C, D). Expectedly, both had frame-shift mutations.

### Assessment of the development of the newly generated Ets2 mutant mice

Previous reports indicated that *Ets2^db1/db1^* mice exhibit an embryonic lethal phenotype due to a placental deficiency (Yamamoto et al., 1998). Sixteen pups were obtained from three derivations, and as expected, none of the pups included the double mutant alleles (*em1/em2*) (Table S1). Further, we analyzed the developmental ability of *Ets2* mutant mice by performing a test cross using *Ets2^+/em1^* mice and assessed the genotypes of the offspring. No homozygous mutant pups (*Ets2^em1/em1^*) were born (wild: hetero: homo = 45: 91: 0, Table S1).

A previous study reported that *Ets2^db1/db1^* embryos were degenerated by the placental deficiency around E7.5 and disappeared after E8.5. To investigate whether the *Ets2^em1/em1^* mutant phenocopies the *Ets2^db1/db1^* mutant, we crossed *Ets2^+/em1^* animals and observed embryos at several stages. *Ets2^em1/em1^* embryos at E7.5 were slightly delayed in their developmental stage but clearly progressed in a comparable manner to embryos from *Ets2^db1/db1^* animals. The *Ets2^em1/em1^* embryos had survived at E8.5, but all of them were retarded. By E9.5 and E10.5, some malformed *Ets2^em1/em1^* embryos were present and developed before the turning of the embryo, which usually occurred at approximately E8.5 (Fig. 2A and Table S2).

**Fig. 2.**
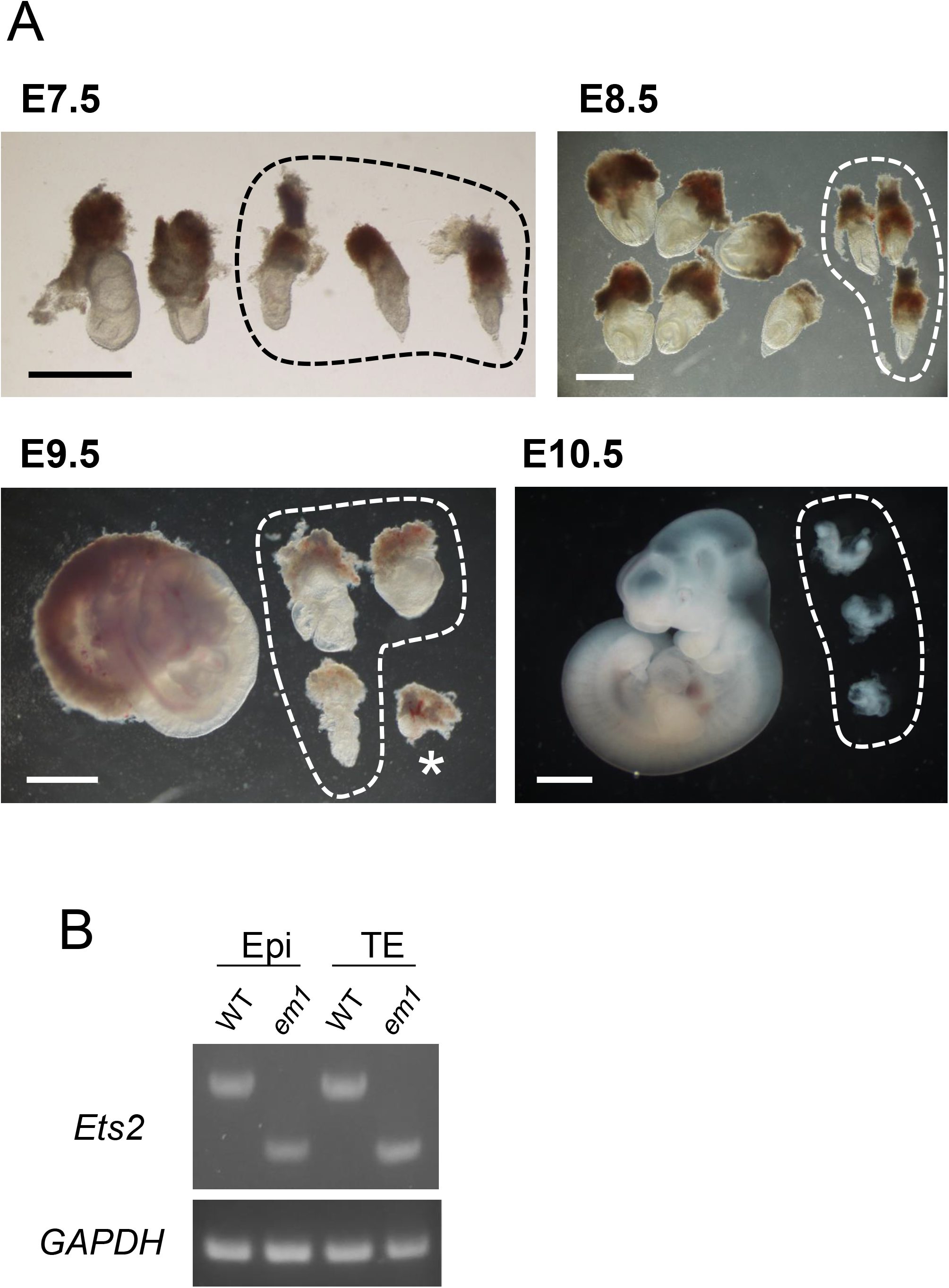
Development of *Ets2 ^em1/em1^* embryos. (A) From E7.5 to E10.5, embryos were observed after crossing *Ets2^+/em1^* females and males. *Ets2^em1/em1^* embryos are indicated as circles in each picture. The regions outside of the circles correspond to *Ets2^+/+^* or *Ets2^+/em1^* embryos. The genotype of the embryo indicated by an asterisk at E9.5 could not be determined. All scale bars indicate 1 mm. (B) Gene expressions of E7.5 WT and *Ets2^em1/em1^* (em1) embryos. Embryos were separated into trophectodermal tissue (TE), including ectoplacental corn, and epiblast (Epi), from which RNA and cDNA were prepared. Both em1 bands were shifted to be lower than the WT bands. The DNA sequences of the em1 bands were the same as em1L in Fig. 4B.

As the frame-shift mutation in *Ets2^em1/em1^* is located in exon 8, the stop codon occurs before exon 9, and the original stop codon is located in exon 10. For this reason, NMD might occur, such that the *em1* mutant mRNA may be degraded in *Ets2^em1/em1^* embryos. To confirm this, we performed RT-PCR using E7.5 embryos. Embryos were separated into the posterior trophectoderm (TE) and anterior epiblast (Epi). Both regions expressed *em1* mutant mRNA, and their sequences included 205 nt deletions that were predicted from the genomic sequence. It is likely the case that the *em1* mutant exhibited a distinct phenotype compared with the Ets2 mutant if the *em1* mRNA was translated into a protein product (Fig. S3).

### Establishment of Ets2 homozygous mutant ESC lines and phenotypic analysis after birth

By rescuing placental function using the tetraploid complementation method, *Ets2^db1/db1^* offspring were successfully developed to term. Therefore, we attempted the same experiment to define whether the embryonic lethal phenotype of *Ets2^em1/em1^* was dependent on the placental deficiency or not.

Before conducting the tetraploid complementation, we established *Ets2^em1/em1^* and *Ets2^em2/em2^* ESC lines. In this way, we improved the efficiency of obtaining homozygous mutant mice because the ratio of homozygous mutant embryos was only one out of four when we used embryos from a heterozygous crossing. After crossing heterozygotes, two-cell embryos were collected and developed until the blastocyst stage. ESC lines were established from the collected blastocysts and analyzed by genotyping. The rate of homozygous mutant ESC line establishment for both the *Ets2^em1/em1^* and *Ets2^em2/em2^* mutants followed Mendel’s law (Table S3).

Using the tetraploid complementation method, offspring were obtained from *Ets2^em1/em1^* and *Ets2^em2/em2^* ESC lines (Figs. 3A, B and Table S4). This result indicated that the embryonic lethality observed for the *Ets2^em1/em1^* and *Ets2^em2/em2^* genotypes was due to a dysfunction of placental differentiation, the same as that seen for the *Ets2^db1/db1^* mutant. Unexpectedly, wavy hair and curly whisker phenotypes were not observed in *Ets2^em1/em1^* or *Ets2^em2/em2^* mice (Figs. 3B, C).

**Fig. 3.**
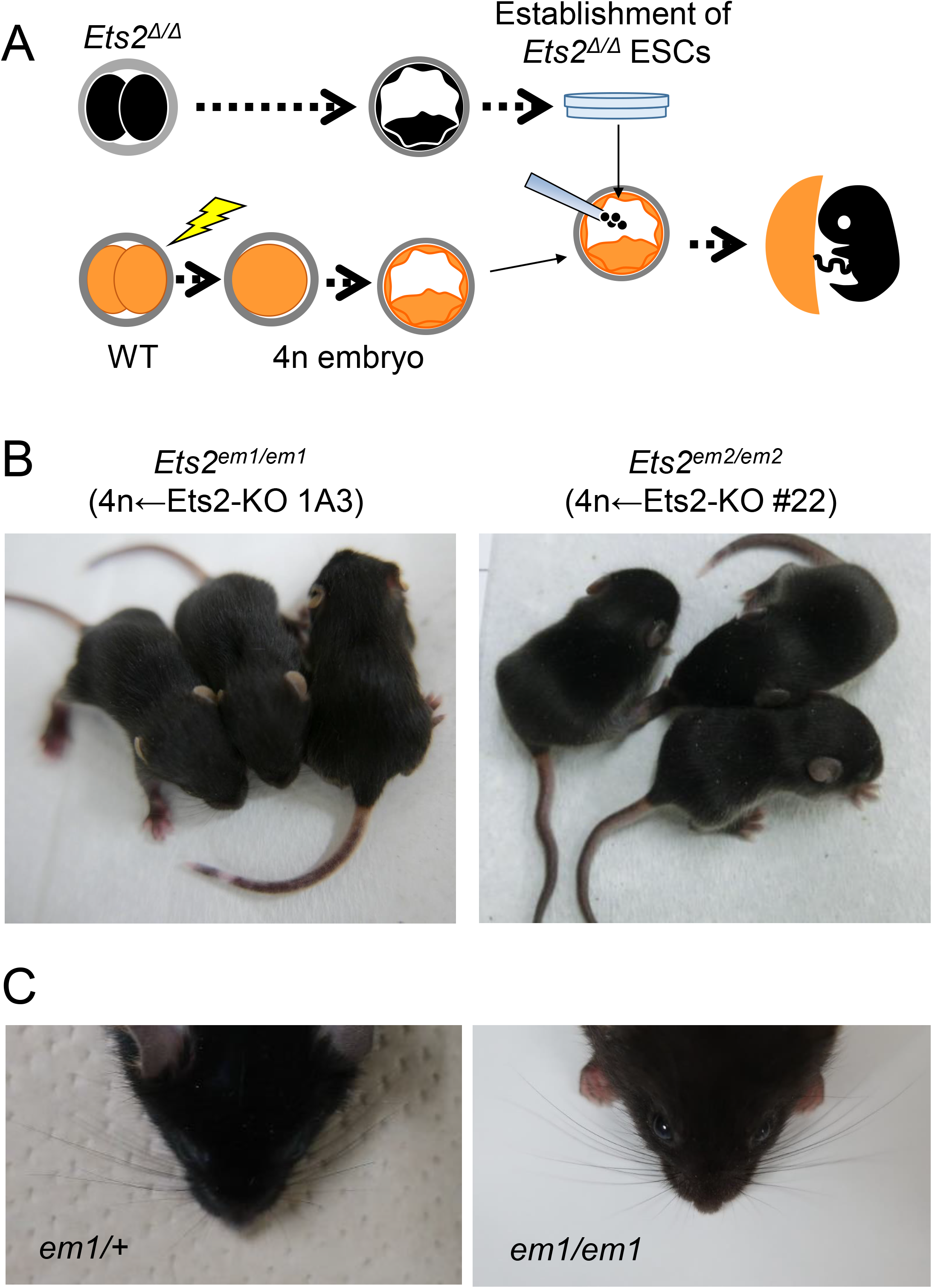
Assessment of hair and whisker phenotypes after birth in newly generated *Ets2* mutant mice. (A) Strategy for the generation of *Ets2* homozygous mutant mouse with ESCs using the tetraploid complementation. (B) 2-week-old *Ets2^em1/em1^* (left picture) and *Ets2^em2/em2^* (right picture) mice. (C) Faces of 4-week-old in *Ets2^+/em1^* (left picture) and *Ets2^em1/em1^* (right picture). Curly whiskers were not observed in *Ets2^em1/em1^*.

To corroborate the relationship between Ets2 and the wavy hair phenotype, we established a null mutant ES cell line, in which a region upstream of exon 2 through the 3’-UTR of exon 10 was deleted, including all open reading frame (ORF) regions. Pups were then generated using the tetraploid complementation method (Fig. S1 and Table S5). Both the wavy hair and curly whisker phenotypes were observed in *Ets2 null* mice from around 2-weeks of age, as was observed for *Ets2^db1/db1^* mice.

### Gene expression in *Ets2^em1/em1^* skin

In this study, newly established *Ets2^em1/em1^* and *Ets2^em2/em2^* mice exhibited an embryonic lethal phenotype due to placental dysfunction but did not exhibit the wavy hair phenotype of *Ets2^db1/db1^* mice, despite having a frame-shift mutation. Therefore, we next investigated the *Ets2* gene expression from the *Ets2* locus in the skin of *Ets2^em1/em1^* mice.

In 4-week-old mice, the expression of the mRNA was detected in the skin of wildtype and *Ets2^em1/em1^* mice but not in the skin of *Ets2 null* mice. Notably, two sizes of fragments were detected in *Ets2^em1/em1^* skin samples (Fig. 4A).

**Fig. 4.**
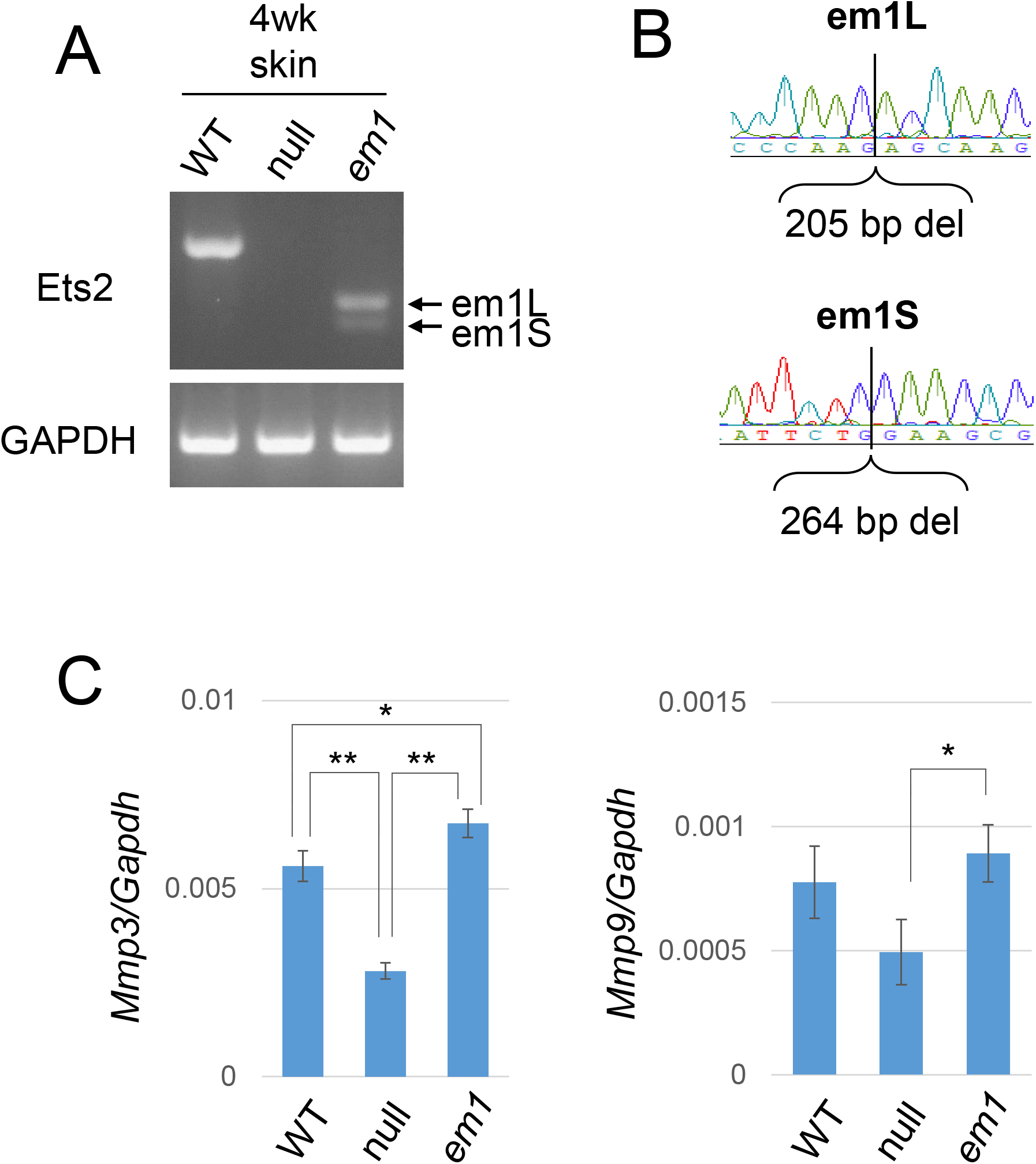
Gene expression in the *Ets2^em1/em1^* skin. (A) Gene expression from the *Ets2* locus in 4-week-old skin of WT, *Ets2 null* mutant (null), and *Ets2 ^em1/em1^* (em1) mice. From em1 skin, two types of mRNA were expressed, although both were shorter than WT. The larger mRNA from em1 skin was called em1L, and the smaller mRNA was called em1S. (B) Sequences of em1L and em1S. Em1L was the expected sequence, but em1S contained a deleted locus that matched exon 8, shown in Fig. S2. (C) Gene expression of *Mmp3* and *Mmp9* in the *Ets2 ^em1/em1^* skin. There were significant differences in the expression level of *Mmp3* between WT and null, em1 and null, and WT and em1. The expression level of *Mmp9* in the null skin was slightly decreased compared with WT, but not significantly, and em1 showed a significant difference compared with null. *p < 0.05, **p < 0.01.

We hypothesized that the unexpected fragment size might be attributed to a splice variant and that this could explain the differences in phenotype between *Ets2^em1/em1^* and *Ets2^db1/db1^* animals. Therefore, we next analyzed the sequences of the potential splice variants. Although the large bands observed for the *Ets2^em1/em1^* skin and thymus and *Ets2^em2/em2^* thymus represented the expected frame-shifted sequences, the sequences of the smaller bands showed skipping of exon 8, which was in-frame and consisted of 264 bp (Fig. 4B and Fig. S2). This Ets2 protein, which skipped exon 8, was predicted to contain the ETS domain based on the SMART online database (SMART) (Fig. S3).

Further, we examined the gene expression of *MMP-3* and *MMP-9* in *Ets2^em1/em1^* skins, since a previous report showed that expression of these genes was decreased in *Ets2^db1/db1^* mice (Yamamoto et al., 1998). However, the expression levels of *MMP-3* and *MMP-9* in 4-week-old *Ets2^em1/em1^* skins were not reduced compared with the wildtype, even though they were reduced in the *Ets2 null* (Fig. 4C).

## Discussion

In this work, we newly generated two *Ets2* mutant models, namely, *em1* and *em2*, which contained frame-shift genomic mutations with the stop codon located before the ETS domain. We predicted that both mutant proteins would lack the ETS domain if they were translated. Therefore, we expected that the phenotypes in these homozygous mutants would mimic those described in previous reports for *Ets2^db1/db1^* mice. Indeed, *Ets2^em1/em1^* mice exhibited the same embryonic lethal phenotype, although the survival period was a little longer than that observed for *Ets2^db1/db1^* mice (Yamamoto et al., 1998).

A previous study suggested that different phenotypes observed in genomic mutants in mice could be attributed to differences in strain backgrounds (Coley et al., 2016; Desroches-Castan et al., 2019; Montagutelli, 2000). The *Ets2^db1/db1^* mutant was established using the Swiss Black and 129/Sv strains of mice, and we established the *Ets2^em1/em1^* and *Ets2^em2/em2^* mutants using the B6D2F1 mix background. The variation between these strains might have resulted in the minor difference in the embryonic lethal phenotype that we observed, which was caused by a trophectodermal abnormality.

On the other hand, the wavy hair and curly whisker phenotypes observed in *Ets2^db1/db1^* mice did not occur in our *Ets2^em1/em1^* and *Ets2^em2/em2^* mice. We found that the skin of *Ets2^em1/em1^* mice expressed a mutant mRNA lacking exon 8 that could potentially be translated into a protein, including the ETS domain. Indeed, it is possible that this exon 8 skip protein, including the ETS domain, could rescue Ets2 function in hair and whisker. This was strongly suggested by the finding that these mice exhibited comparable levels of *MMP-3* and *MMP-9* mRNAs to wildtype, and *Ets2^db1db1^* and *Ets2^db1db1^* mice exhibited decreased expression.

DNA mutations in a genetic locus frequently lead to exon skipping, and several human diseases are linked to these types of exon skipping events. In addition, exon skipping can occur because of lack of exonic splicing enhancer sequences or an exonic splicing silencer sequence inside of an exon (Baralle and Giudice, 2017; Cartegni et al., 2002).

Some reports have also suggested that unexpected exon skips can occur when using the CRISPR/Cas9 system, indicating that a frame-shifted mutant exon induced by CRISPR/Cas9 is skipped but can be induced through alternative splicing or in-frame exon skipping (Chen et al., 2018; Mou et al., 2017; Sui et al., 2018; Tuladhar et al., 2019). Since the exon 8 skip mRNA in the skin and thymus of *Ets2^em1/em1^* is in-frame and the *em1* mRNA sequence was designed as a frame-shift mutation, we believe that a portion of the *em1* pre-mRNA could have been spliced through the same mechanism as described in previous studies (Chen et al., 2018; Mou et al., 2017; Sui et al., 2018; Tuladhar et al., 2019).

Further, in this study, the exon skip found was only identified in the skin and thymus and was not detected during embryonic stages. Thus, the induction of the exon skip was likely due to the changes in mRNA splicing that were dependent on the cell type or developmental stage, despite having the same genomic mutation. This phenomenon has a possible effect on the appearance of phenotypes. This is a novel finding of this study. Further, this finding suggested that all ORF-deletion models are adequate for analysis of the genes’ functions.

During the development of the neuron and heart, tissue-specific RNA binding proteins that induce alternative splicing are expressed (Baralle and Giudice, 2017). Moreover, epigenetic modifications, such as DNA methylation and histone modifications, can result in tissue-specific alternative splicing (Baralle and Giudice, 2017). Thus, these phenomena might affect the differences in the splicing pattern in the mutant tissues.

Ets2 plays a role not only in trophoblast formation and hair morphology but also in cancer, angiogenesis, and the immune system. Therefore, the newly generated *Ets2* mutant models in this report will likely be applicable to several different research fields, such as physiological research *in vivo* and molecular biological research into splicing.

## Methods

### Animals

All animal experiments were conducted in accordance with the guidelines of “Regulations and By-Laws of Animal Experimentation at the Nara Institute for Science and Technology,” and were approved by the Animal experimental Committee at the Nara Institute of Science and Technology (the approval no.1639). B6D2F1 female mice and ICR mice were purchased from SLC (Japan). C57BL/6J male mice were purchased from CLEA (Japan).

### Collection of zygotes

Female mice were treated by PMSG and hCG for superovulation, then mated with male mice. Pronuclear stage zygotes were collected from female oviducts after 20 hours of hCG injection. After removing cumulus cells using hyaluronidase, zygotes were incubated in KSOM at 37°C under 5% CO2 in the air until use. 2-cell stage zygotes were collected from female oviducts after 42-46 hours of hCG injection by the flush-out method. Collected 2-cell stage embryos were incubated until use the same as above.

### Generation of *Ets2* mutant zygote by CRISPR/Cas9 system using electroporation

Target sites of guide RNA (gRNA) were designed using the web tool CRISPR direct [24]. Genome editing by electroporation was performed as a previous study [12].

CFB16-HB and LF501PT1-10 electrode (BEXCo.Ltd., Tokyo, Japan) were used for electroporation. 30–40 pronuclear stage zygotes were subjected to electroporation at one time. Zygotes were washed with Opti-MEM I (Thermofisher) three times, subsequently placed in a line in the electrode gap filled with 5 μl the mixture of 120 ng/μl Cas9 protein (TaKaRa, Japan), 300 ng/μl *tracerRNA,* and 200 ng/μl *crRNA* (HPLC grade, Fasmac) in Opti-MEM I. The electroporation condition was performed were 30V (3 msec ON ± 97 msec OFF) four times. After electroporation, zygotes were washed with KSOM three times then cultured until developing the eight-cell stage. Eight-cell stage embryos were provided to the tetraploid complementation.

### Establishment of *Ets2* mutated ESC lines

To establish *the Ets2* homozygous mutant model, collected 2-cell stage embryos from *Ets2* heterozygous mutant parents were incubated until the blastocyst stage, removing the Zona pellucida (ZP) using Acidic Tyrode solution (Sigma T1788). Blastocyst embryos without the ZP were seed on gelatin-coated 60-mm dishes and cultured on mouse embryonic fibroblast (MEF) with N2B27 medium supplemented with 3 μM CHIR99021(Axon1386), 1.5 μM CGP77675 (Sigma SML0314), and mouse LIF (N2B27-a2i/L medium) [25]. After seven days, the outgrowth of blastocysts was disaggregated by 0.25% trypsin in 1mM EDTA in PBS (-). Half of the cells were seeded on MEF with the gelatin-coated dishes for expanding. The others were seeded on the gelatin-coated dishes without MEF for genotyping by PCR. *Ets2* homozygous mutant ESC lines were provided for tetraploid complementation. The *Ets2* null mutant model was established using ESCs, as performed in a previous study (Oji et al., 2016). mF1-05 ESC line, which was newly established from 129X1and C57BL6/J F1 embryo, was seeded on MEF then transfected with two designed pSpCas9(BB)-2A-Puro (pX459) V2.0 (Addgene #62988) plasmids using Lipofectamine 3000 (Thermofisher). Transfected cells were selected by transient treatment with 1 μg/ml puromycin; then, ESC colonies were subject to genotyping with PCR and sequencing. The *Ets2* null mutant ESC line was provided for tetraploid complementation.

### Tetraploid complementation

Tetraploid embryos were prepared as described previously(Kokubu et al., 2009; Okada et al., 2007). In brief, ICR two-cell stage embryos were placed in a fusion buffer, and electrofusion was performed by applying 140 V for 50 ms after aligning embryos between the electrodes. CFB16-HB and LF501PT1-10 electrode (BEXCo.Ltd., Tokyo, Japan) were used for cell fusion.

A wild-type tetraploid four-cell embryo and a genome-edited diploid eight-cell embryo were aggregated after removing the zona pellucida for the aggregation method. For the injection method, *Ets2* mutant ESCs were injected into a wild-type tetraploid four-cell embryo or blastocyst. These embryos were cultured until the blastocysts stage and transferred into the uterus of E2.5 pseudopregnant ICR mice. Offspring were recovered by natural delivery or Caesarean section on E19.5. The mutation of the offspring was detected by genotyping with PCR and sequencing.

### Genotyping

Genotyping primers for detecting *Ets2*-wild, em 1, and em 2 alleles were 5’-ctgagtttaagagtgctcggagg-3’ (Ets2_Fw) and 5’-gccctataggacttgtgtacagg-3’ (Ets2_Rev). Primers for Ets2 null mutant allele(s) were 5’-tgtggagtctcacatcgaag-3’ (Ets2_Ex2_F) and 5’-gggcctgctcggtgccacgg-3’ (Ets2_EX10_R). DNA fragments were amplified using GoTaq (Promega) for 40 cycles under the following conditions: 94 °C for 30 sec, 60 °C for 30 sec and 68 °C for 40 sec for detecting wild, em1 or em2 allele, and 94 °C for 30 sec, 60 °C for 30 sec and 68 °C for 20 sec for detecting the null allele, respectively.

### RNA expression analysis

Mouse cDNAs were prepared from 4-week old skin, adult skin, and adult thymus using SuperScript III Reverse Transcriptase (Thermo Fisher Scientific) after purified RNA by Trizol reagent (Thermo Fisher Scientific). RT-PCR was performed using 20 ng of cDNA with the following primers: 5’-CGTGAATTTGCTCAACAACAATTCTG-3’ and 5’-gagaggctatgccggt-3’ for *Ets2* and 5’-CCAGTATGACTCCACTCACG-3’ and 5’-GACTCCACGACATACTCAGC-3 for *Gapdh* (Wen et al., 2007). cDNA fragments were amplified using KOD Fx Neo (TOYOBO) or GoTaq (Promega) for 35 cycles under the following conditions: 94 °C for 30 sec, 60 °C for 30 sec and 72 °C for 40 sec for *Ets2,* and 94 °C for 30 sec, 53 °C for 30 sec and 72 °C for 30 sec for *Gapdh.*

Quantitative real-time PCR was performed using 20 ng of cDNA with following primers: 5’-TTAAAGACAGGCACTTTTGG-3’ and 5’-CAGGGTGTGAATGCTTTTAG-3’for *Mmp3,* 5’-CGTCTGAGAATTGAATCAGC-3’ and 5’-AGTAGGGGCAACTGAATACC-3’ for *Mmp9* expression (Man et al., 2003). Gene expression level was normalized by Gapdh, the same cDNA. The primer set for *Gapdh* was the same as above. Real-time PCR was performed by LightCycler96 (Roche) using Luna Universal qPCR Master Mix (NEB), and the data were analyzed by LightCycler software.

### Statistics analysis

The statistical difference was determined using the Student t-test. Differences were considered statistically significant if the P-value was less than 0.05.

## Acknowledgements

The authors would like to thank Enago (www.enago.jp) for the English language review.

## Competing interests

The authors declare no competing financial interests.

## Author contributions

A.I., Y.K., I.N. performed most experiments, assisted by N.Y. who performed tetraploid complementation, and S.Y., performed qPCR. A.I., S.Y., and M.I. analyzed the data. A.I. wrote the manuscript and all authors discussed the results and commented on the manuscript.

## Funding

This work was supported by JSPS KAKENHI Grant Number 16K07091, 18H04885, Start Up Fund for female researchers in NAIST, and KAC 40th Anniversary Research Grant.

## References

Baralle, F. E. and Giudice, J. (2017). Alternative splicing as a regulator of development and tissue identity. Nat Rev Mol Cell Biol 18, 437–451.

Baran, C. P., Fischer, S. N., Nuovo, G. J., Kabbout, M. N., Hitchcock, C. L., Bringardner, B. D., McMaken, S., Newland, C. A., Cantemir-Stone, C. Z., Phillips, G. S., et al. (2011). Transcription factor ets-2 plays an important role in the pathogenesis of pulmonary fibrosis. Am JRespir Cell Mol Biol 45, 999–1006.

Cartegni, L., Chew, S. L. and Krainer, A. R. (2002). Listening to silence and understanding nonsense: exonic mutations that affect splicing. Nat Rev Genet 3, 285–298.

Chen, D., Tang, J. X., Li, B., Hou, L., Wang, X. and Kang, L. (2018). CRISPR/Cas9-mediated genome editing induces exon skipping by complete or stochastic altering splicing in the migratory locust. BMC Biotechnol 18, 60.

Coley, W. D., Bogdanik, L., Vila, M. C., Yu, Q., Van Der Meulen, J. H., Rayavarapu, S., Novak, J. S., Nearing, M., Quinn, J. L., Saunders, A., et al. (2016). Effect of genetic background on the dystrophic phenotype in mdx mice. Hum Mol Genet 25, 130–145.

Cong, L., Ran, F. A., Cox, D., Lin, S., Barretto, R., Habib, N., Hsu, P. D., Wu, X., Jiang, W., Marraffini, L. A., et al. (2013). Multiplex genome engineering using CRISPR/Cas systems. Science 339, 819–823.

Desroches-Castan, A., Tillet, E., Ricard, N., Ouarné, M., Mallet, C., Feige, J. J. and Bailly, S. (2019). Differential Consequences of. Cells 8.

Hashimoto, M., Yamashita, Y. and Takemoto, T. (2016). Electroporation of Cas9 protein/sgRNA into early pronuclear zygotes generates non-mosaic mutants in the mouse. Dev Biol 418, 1–9.

Karim, F. D., Urness, L. D., Thummel, C. S., Klemsz, M. J., McKercher, S. R., Celada, A., Van Beveren, C., Maki, R. A., Gunther, C. V. and Nye, J. A. (1990). The ETS-domain: a new DNA-binding motif that recognizes a purine-rich core DNA sequence. Genes Dev 4, 1451–1453.

Kokubu, C., Horie, K., Abe, K., Ikeda, R., Mizuno, S., Uno, Y., Ogiwara, S., Ohtsuka, M., Isotani, A., Okabe, M., et al. (2009). A transposon-based chromosomal engineering method to survey a large cis-regulatory landscape in mice. Nat Genet 41, 946–952.

Mali, P., Yang, L., Esvelt, K. M., Aach, J., Guell, M., DiCarlo, J. E., Norville, J. E. and Church, G. M. (2013). RNA-guided human genome engineering via Cas9. Science 339, 823–826.

Man, A. K., Young, L. J., Tynan, J. A., Lesperance, J., Egeblad, M., Werb, Z., Hauser, C. A., Muller, W. J., Cardiff, R. D. and Oshima, R. G. (2003). Ets2-dependent stromal regulation of mouse mammary tumors. Mol Cell Biol 23, 8614–8625.

Montagutelli, X. (2000). Effect of the genetic background on the phenotype of mouse mutations. J Am Soc Nephrol 11 Suppl 16, S101–105.

Mou, H., Smith, J. L., Peng, L., Yin, H., Moore, J., Zhang, X. O., Song, C. Q., Sheel, A., Wu, Q., Ozata, D. M., et al. (2017). CRISPR/Cas9-mediated genome editing induces exon skipping by alternative splicing or exon deletion. Genome Biol 18, 108.

Nagy, A., Rossant, J., Nagy, R., Abramow-Newerly, W. and Roder, J. C. (1993). Derivation of completely cell culture-derived mice from early-passage embryonic stem cells. Proc Natl Acad Sci USA 90, 8424–8428.

Oji, A., Noda, T., Fujihara, Y., Miyata, H., Kim, Y. J., Muto, M., Nozawa, K., Matsumura, T., Isotani, A. and Ikawa, M. (2016). CRISPR/Cas9 mediated genome editing in ES cells and its application for chimeric analysis in mice. Sci Rep 6, 31666.

Okada, Y., Ueshin, Y., Isotani, A., Saito-Fujita, T., Nakashima, H., Kimura, K., Mizoguchi, A., Oh-Hora, M., Mori, Y., Ogata, M., et al. (2007). Complementation of placental defects and embryonic lethality by trophoblast-specific lentiviral gene transfer. Nat Biotechnol 25, 233–237.

Seidel, J. J. and Graves, B. J. (2002). An ERK2 docking site in the Pointed domain distinguishes a subset of ETS transcription factors. Genes Dev 16, 127–137.

Sharrocks, A. D. (2001). The ETS-domain transcription factor family. Nat Rev Mol Cell Biol 2, 827–837. SMART http://smart.embl-heidelberg.de/.

Sui, T., Song, Y., Liu, Z., Chen, M., Deng, J., Xu, Y., Lai, L. and Li, Z. (2018). CRISPR-induced exon skipping is dependent on premature termination codon mutations. Genome Biol 19, 164.

Tuladhar, R., Yeu, Y., Tyler Piazza, J., Tan, Z., Rene Clemenceau, J., Wu, X., Barrett, Q., Herbert, J., Mathews, D. H., Kim, J., et al. (2019). CRISPR-Cas9-based mutagenesis frequently provokes on-target mRNA misregulation. Nat Commun 10, 4056.

Wei, G., Guo, J., Doseff, A. I., Kusewitt, D. F., Man, A. K., Oshima, R. G. and Ostrowski, M. C. (2004). Activated Ets2 is required for persistent inflammatory responses in the motheaten viable model. J Immunol 173, 1374–1379.

Wei, G., Srinivasan, R., Cantemir-Stone, C. Z., Sharma, S. M., Santhanam, R., Weinstein, M., Muthusamy, N., Man, A. K., Oshima, R. G., Leone, G., et al. (2009). Ets1 and Ets2 are required for endothelial cell survival during embryonic angiogenesis. Blood 114, 1123–1130.

Wen, F., Tynan, J. A., Cecena, G., Williams, R., Múnera, J., Mavrothalassitis, G. and Oshima, R. G. (2007). Ets2 is required for trophoblast stem cell self-renewal. Dev Biol 312, 284–299.

Yamamoto, H., Flannery, M. L., Kupriyanov, S., Pearce, J., McKercher, S. R., Henkel, G. W., Maki, R. A., Werb, Z. and Oshima, R. G. (1998). Defective trophoblast function in mice with a targeted mutation of Ets2. Genes Dev 12, 1315–1326.

